# Repeated Early Adolescence Ethanol Intoxication Promotes Riskier Decision-Making in Adult Males and Increases Drinking in Adult Female mice

**DOI:** 10.1101/2025.08.03.668331

**Authors:** Leticia Souza Pichinin, Myrna Guernelli, Marianna Nogueira Cecyn, Beatriz Deo Sorigotto, Karina Possa Abrahao

## Abstract

Exceeding binge-level drinking is common among adolescents and is associated with both short- and long-term adverse outcomes. This study evaluated the effects of repeated ethanol-induced intoxication during early adolescence on behavioral outcomes in early adulthood. Male and female C57BL/6J mice (5 weeks old) received intraperitoneal (i.p.) injections of either saline (control) or 3.2 g/kg ethanol (intoxicated). A subset of intoxicated mice had blood collected on days 1 and 4, confirming heavy intoxication (∼289 mg/dL). At 9 weeks of age, animals were tested in the Light-Dark Box (LDB) and Elevated Plus Maze (EPM) immediately after receiving either saline or 1.2 g/kg ethanol. In the LDB, early adolescence intoxication did not affect anxiety-like behaviors but reduced risk assessment behaviors in males, indicating riskier decision-making. No pretest ethanol effect was observed. In the EPM, pretest ethanol produced an anxiolytic effect, accompanied by increased exploration and a reduction in risk-assessment behaviors, while adolescence intoxication did not yield significant effects. To better characterize behavioral organization beyond discrete measures, we applied Markov chain models to quantify first-order transition probabilities to and from risk-assessment states. This analysis revealed that pretest ethanol markedly reduced the complexity of behavioral structure, especially in the EPM, while adolescent intoxication had no detectable ethological effect. Finally, a separate cohort of controls and intoxicated mice underwent a two-bottle choice Intermittent Overnight Drinking protocol in adulthood. Adolescent ethanol intoxication increased voluntary ethanol intake in females but not in males. These findings highlight long-lasting and sex-specific consequences of early adolescent intoxication on risk-related behaviors and alcohol consumption.

## 1. Introduction

Episodic ethanol intoxication increases the risk of injuries and other harmful situations and represents a risk factor for the development of alcohol use disorders (AUD) (1) later in life. In Brazil, 5% of adolescents (12-17 years) and 20.5% of young adults (18-24 years) reported at least one episode of binge drinking in the last 30 days (2). According to the National Institute on Alcohol Abuse and Alcoholism (NIAAA) at the National Institutes of Health (NIH), binge drinking is defined as consuming four to five standard drinks within two hours, typically resulting in blood ethanol concentrations (BECs) above 80 mg/dL. However, high-intensity drinking (Level II: 8–10 drinks; and Level III: 12–15 drinks) can produce even higher BECs, exceeding 200 mg/dL, a threshold considered dangerous by the World Health Organization (3). At these levels, ethanol intoxication impairs motor control and cognitive functions, leading to sedation and, in some cases, loss of consciousness (4). Additionally, it may produce anterograde amnesia (i.e., blackouts) (5,6).

C57BL/6 mice are commonly used as animal models for studying alcohol intoxication due to their high levels of voluntary ethanol consumption. However, animals do not typically drink to the point of reaching BECs above 200 mg/dL or experiencing, in some cases, loss of consciousness (i.e., passing out). Therefore, non-contingent (experimenter-administered) peripheral ethanol administration is necessary to achieve high levels of intoxication. To model high-intensity intoxication, BEC must reach 200 mg/dL or more, which are typically observed at doses above 3.0 g/kg (7). At these doses, mice might lose their righting reflex (LORR), which refers to the animal’s innate ability to correct its body orientation when placed on its back. Notably, even lower doses, such as 2.4 g/kg, have been shown to produce amnestic effects, indicating the impact on memory (8).

Adolescence is a developmental window characterized by heightened behavioral disinhibition, impulsivity, and risk-taking, factors strongly associated with alcohol misuse and escalation of intake (9,10). In addition, alcohol intoxication during adolescence can lead to the persistence of adolescent-like behavioral traits into adulthood (11–15). Converging evidence demonstrates that adolescent ethanol exposure biases individuals toward behavioral phenotypes characterized by poor inhibitory control, maladaptive evaluation of risk, and riskier decision-making (e.g., 16-20). On the other hand, studies examining the long-term consequences of early-adolescent ethanol intoxication on the intricate relationship among anxiety-like, exploratory and risk-assessment behaviors, as well as voluntary drinking in adult mice are still limited (21,22), particularly with regard to sex differences and ethologically informed behavioral measures.

In the present study, we investigated how repeated high-intensity ethanol intoxication during early adolescence affects adult anxiety-like behavior, exploratory strategies, ethological risk-assessment, and voluntary ethanol consumption in female and male C57BL/6 mice. By modeling intoxication levels relevant to human high-intensity drinking, this study aims to clarify how ethanol exposure during a developmental window of heightened vulnerability can shape risk-related behavioral phenotypes into adulthood.

## 2. Methods

### 2.1 Animals

This study was approved by the Comissão de Ética no Uso de Animais (CEUA) of the Universidade Federal de São Paulo (UNIFESP) (#2423170220). The experiments were in accordance with the Conselho Nacional de Controle de Experimentação Animal (CONCEA). Female and male C57BL/6 mice were obtained from Centro de Desenvolvimento de Modelos Experimentais para Biologia e Medicina (CEDEME). Animals arrived at the fourth week of age and were group-housed in automatically ventilated cages (Alesco ALBR Indústria e Comércio Ltda, Monte Mor, Brazil) with 3-4 animals of the same sex per cage. *Vivarium* temperature was kept at 21 ± 2 °C, with 60% humidity and a 12/12-hour light-dark cycle (lights on at 7 a.m.). The animals had *ad libitum* access to standard chow (Nuvilab CR-1 Chow, Nuvital, Paraná, Brazil) and water. Cages contained a half-cardboard roll and a paper sheet, and materials were changed every week. Animals were habituated to the *vivarium* and to the experimenter’s handling before the beginning of the experiments.

All experiments were conducted with female and male mice to allow proper comparison between sexes. All apparatuses (LORR vertices, Elevated Plus Maze – EPM, and Light–Dark Box – LDB) were cleaned with 70% ethanol solution and allowed to fully evaporate and dry before placing the animals. Anxiety-like behavior tests and ethanol intake procedures were initiated during the 9^th^ week of life, corresponding to early adulthood (10). The experimental design for the three experiments conducted in this study is illustrated in Figure 1.

**Figure 1:**
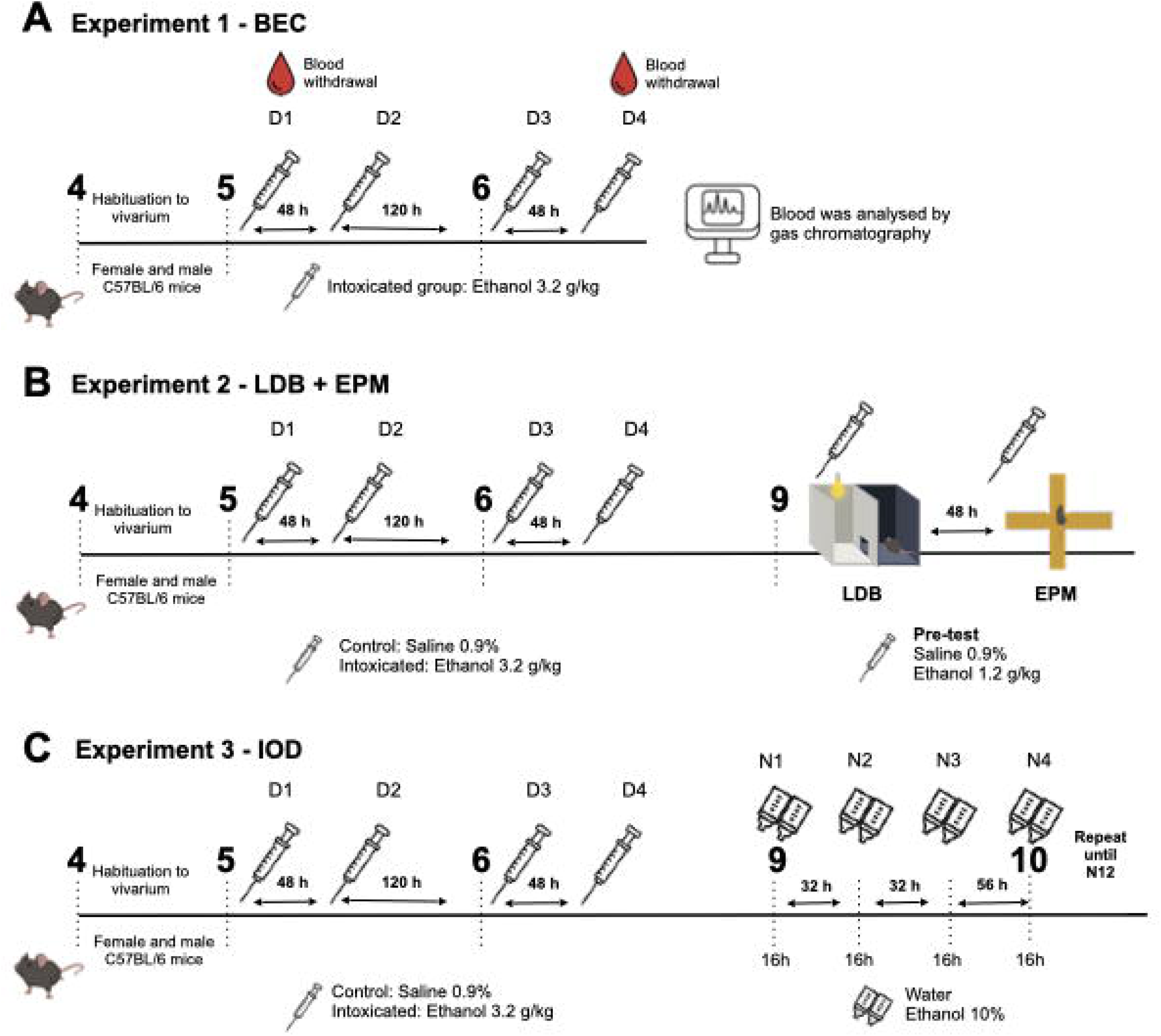
Experimental design. All experiments were conducted in female and male C57BL/6 mice. The timeline depicts the animals’ age in weeks and the interval in hours between procedures. Syringe icons represent intraperitoneal (i.p.) injections, and bottles icons represent access to 10% ethanol and water. “D” stands for day and “N” stands for night. **(A)** Experiment 1 -BEC. A single group of mice was intoxicated (3.2 g/kg, i.p.) during LORR on four occasions (D1–D4), and blood samples were collected on D1 and D4 for blood ethanol concentration (BEC) analysis by gas chromatography. **(B)** Experiment 2 – LDB + EPM. Mice were assigned to either an intoxicated group (3.2 g/kg, i.p.) or a control group (0.9% saline) during LORR on D1–D4. In week 9, animals underwent the Light–Dark Box (LDB) test, followed 48 h later by the Elevated Plus Maze (EPM). Before each behavioral test, mice received an acute injection of ethanol (1.2 g/kg) or saline, generating four experimental conditions: intoxicated–ethanol, intoxicated–saline, control–ethanol, and control–saline. **(C)** Experiment 3 – IOD. Mice were assigned to intoxicated (3.2 g/kg, i.p.) or control (0.9% saline) groups during LORR on D1–D4. In week 9, all animals began the intermittent ethanol access protocol (Intermittent Open Drinking, IOD), consisting of alternating nights with 10% ethanol and water available, following standard IOD cycles (N1–N12).

### 2.2 Ethanol-induced loss of righting reflex (LORR)

To model high levels of intoxication, we used the ethanol-induced LORR protocol. One day prior to the first intoxication, animals were habituated to an intraperitoneal injection (i.p.) of saline (0.9% NaCl; 4.0 ml/kg, matching the ethanol injection volume). LORR protocol consisted of four ethanol administrations during early adolescence (postnatal weeks 5-6): the first two injections were spaced by 48 hours, followed by a third injection five days later and a fourth injection 48 hours after that. Mice received saline (0.9% w/v) (control group) or 3.2 g/kg of ethanol (20% v/v, prepared from 95% ethanol; LabSynth, São Paulo, Brazil) (intoxicated group) via i.p administration and were immediately placed in acrylic vertices chambers (USA Acrylics, Sao Paulo, Brazil) for monitoring. We selected the 3.2 g/kg ethanol dose because it induces robust behavioral effects and results in blood ethanol concentrations (∼289 mg/dL) consistent with NIH and WHO criteria for heavy intoxication. Loss of reflex was defined as the inability of the animal to return to an upright position within three minutes after injection. Mice who failed to lose reflex were placed on warm pads and had data excluded from the LORR analysis on that day specifically (23). Mice were considered to have recovered the reflex when able to turn into the supine position three times in a row within one minute. After recovering the reflex, animals were taken back into their home cages and left on warm pads for at least one hour to allow complete recovery. Wet chow food was offered, and saline injections were administered to animals in case they weren’t moving and/or eating normally one hour after the procedure. Animals in the control group were also placed on warm pads for one hour. It is important to note that all mice, including those that did not exhibit LORR after ethanol administration, were included in the final behavioral analyses. Similar results were observed when analyses were conducted exclusively on mice that exhibited loss of the righting reflex (data not shown).

### 2.3 Blood Ethanol Concentration

A group of mice (Experiment 1, N = 7-8/sex, Figure 1A) went through the ethanol-induced LORR protocol and had their blood samples collected on days 1 and 4 to assess BECs right on the recovery time. Tail blood was collected in cooled EDTA K2 tubes and briefly vortexed and then frozen at −20 °C. Blood samples were stored until analysis by gas chromatography (Shimadzu, Barueri, Brazil) at CIATox Lab of the Universidade Estadual de Campinas (Unicamp). The curve had 10 standard points, and detection ranged from 0.05 to 5.00 g/L. This experiment was conducted in a single iteration, and all animals were euthanized after the second blood collection.

### 2.4 Light-Dark Box (LDB) and Elevated Plus Maze (EPM)

Another group of mice (Experiment 2, N = 10-13/group/sex, Figure 1B) underwent the ethanol-induced LORR protocol and, in the 9th week, were submitted to the LDB test and, 48h later, to the EPM. This experiment was done in four iterations. Although some differences in the frequency with which specific behaviors were recorded, the overall behavioral trends remained consistent among iterations. Therefore, we opted to compile data from all iterations to ensure sufficient sample size per group and increase the statistical power of the analysis. Half of the animals in each previously described group (intoxicated or control) received i.p. saline (0.9% w/v) or 1.2 g/kg of ethanol (15% v/v), resulting in four groups: control-saline, control-ethanol, intoxicated-saline, and intoxicated-ethanol. Animals were placed in the LDB (in the dark compartment) and the EPM (facing the open arm) one minute after the i.p. administration. Each test lasted five minutes. The 1.2 g/kg ethanol dose was selected based on prior evidence showing its ability to acutely reduce anxiety-like behavior in C57BL/6J mice (24).

The LDB apparatus had two equal-sized (24 cm x 24 cm) compartments with flat floors. The light side (white colored) had approximately 400 lux, and the dark side (black colored), 3 lux. After injection mice were placed in the dark side and the door between compartments was lifted 5 s later. Animal exploration of the light side was recorded with a Digital Camera (Sony Corporation, Japan). One animal did not make the transition to the light side and was excluded from the analysis. The EPM consisted of a wood apparatus elevated 50 cm from the ground with two open arms and two closed arms, with all arms measuring 50 cm. The open arms had an average light of 46 lux, and the closed arms 16 lux. The apparatus was recorded using Ethovision Noldus XT6 (Noldus Information Technology, Netherlands).

Definitions of behaviors analyzed in each test are listed in Supplementary Table 1. Behavior variables such as time spent in each compartment or armside, latency to the first light side entry (for the LDB), number of transitions, head out, stretch, and rearing, sniffing, head dips (for the EPM) were assessed manually with the BORIS software (25) (https://www.boris.unito.it/), with the experimenters blind to the treatment groups. We calculated the “total risk assessment” as the head out and stretch units sum (head outs + stretches). In addition to these measures, we also categorized risk assessment behaviors into NoGo (when animals assessed but did not enter the aversive compartment) and Go (when animals explored the aversive compartment after assessment), to better capture decision-making under approach-avoidance conflict.

### 2.5 Two-bottle choice Intermittent Overnight Drinking (IOD) protocol

A new cohort of mice (Experiment 3; N = 17–18/group/sex; Figure 1C) underwent the ethanol-induced LORR protocol and were later exposed to the IOD protocol beginning in the 9th week of life. This experiment was conducted across three separate iterations, and data from all iterations were pooled to ensure adequate sample size per group and to increase statistical power. For each drinking session, mice were placed individually in cages for 16 hours (starting two hours before lights off and ending two hours after lights on) with *ad libitum* access to food and two bottles (25 mL conical tubes) containing water. The IOD protocol consisted of 12 alternating overnight sessions, with no more than three sessions per week, in which mice were individually housed and provided access to one bottle containing water and the another containing 10% ethanol (26). The positions of the bottles were alternated on each drinking night to prevent side bias. Although housed individually, mice could still see and smell each other. All bottles were weighed before and after each drinking session, and an additional reference bottle was used to correct for leakage and spillage.

### 2.6 Statistical Analysis

We used Generalized Estimated Equations (GEE) with Auto-Regressive (AR1) correlation matrix and Gamma model type, with animal ID as subject variable and day (time) as within-subject variable for LORR and BEC data. Latency to lose the reflex and the duration of LORR were both analyzed in minutes. Controlling variables were experiment iteration, day, and sex. All comparisons were made to day one of intoxication.

For the LDB test, six animals were excluded due to recording issues: three females (one from each group except control-saline) and three males (one from each group except intoxicated-ethanol). For the EPM test, three animals were excluded for the same reason: two males (one from the control–saline group and one from the intoxicated–saline group) and one female (from the intoxicated–saline group). Additionally, animals whose time spent in the light side of the LDB or in the open arms of the EPM fell outside the range defined as mean ± 2 standard deviations (SD) of their respective groups were considered outliers. Based on this criterion, five males were excluded from the LDB dataset: one from the control-saline group, one from the control-ethanol group, one from the intoxicated-saline group, and two from the intoxicated-ethanol group. For the EPM test, one male from the intoxicated–ethanol group was excluded as an outlier. One female from the control–ethanol group was also removed from the EPM dataset because she remained immobile throughout the entire test. Finally, Generalized Linear Models (GzLM) were applied to analyze LDB and EPM datasets, with Linear distribution for continuous numerical variables and Poisson distribution for count data (as number of stretches, for example), both with Log link function. The analyses used intoxication (saline or ethanol during early adolescence), pretest (saline or ethanol right before each behavioral test) and sex as predictor variables.

The exported data from Boris was pre-processed using the App for Behavior Data Processing 3.0, developed in Python (3.11) with Anaconda Environment (https://github.com/mariannacecyn/App-for-Behavior-Data-Processing.git.). To understand how behaviors occur and the relationships between them, we used the Markov Model to study behavior sequence complexity as stochastic events (27,28). First, we calculated the probabilities of each behavior occurring, the initial state vector of MM. Second, we also constructed the Markov Matrix with transition probabilities from the possible transitions with each origin (start) behavior. Then the probability of each transition was calculated as conditional probability, which means mathematically as P(A|B) = p(B)*p(B->A), or in other words, probability of transition B to A is probability of B occurring times the probability of “if in B in time t, A occurs in time t+1”. Since the literature on how to compare groups in this model is still scarce (29,30), we compared the probability of each transition among groups rather than the entire stochastic system. Statistical significance (p < 0.05) is marked as asterisks on the transition vector of the Markov Model graph. Our analysis focused on transitions “to” and “from” aversive compartments. To construct the graphical schema of Markov Model graphs into figures, we separated the transition matrices and state vectors into groups and calculated the mean (≥ 0.02) and SD.

Data from IOD protocol was checked for outliers, defined as values deviating from the group mean by more than ± 2 SD. We used this cutoff as it allowed us to identify procedural issues such as leakage and clogging of bottles. Animals identified as outliers for at least 6 nights were excluded from the dataset. A total of five animals were removed based on this criterion. For intake data, although the Gamma model had a better fit compared to the Linear model, water consumption dataset had 38% zeros, which would be excluded from Gamma analysis. Therefore, we used the GEE Linear Model. The model was built similarly to that described for LORR.

All data were analyzed using JAMOVI 2.4.11 or IBM SPSS Statistics 21. All graphs were created in GraphPad Prism (Version 9.5.1 - 733). Alpha levels were 0.05. When appropriate, the adjusted Bonferroni *post hoc* test was applied to identify specific differences among groups. Data are presented in graphs as mean and standard error means (SEM). Results present the Wald Chi-Square (X²) values, degrees of freedom, and p-value when significant. When necessary, values from parameter estimates and *post hoc* are also shown.

## 3. Results

### 3.1 Experiment 1: High BEC confirming heavy intoxication

Not all mice lost the righting reflex following ethanol administration: 25% did not lose the reflex on day 1, 37% on day 2, 30% on day 3, and 31% on day 4. These rates highlight interindividual variability in sensitivity to ethanol-induced sedation. For the subset of mice that did lose the reflex, we analyzed the latency to LORR in Experiment 1 (BEC) and found no significant effect of day (X²(_([0-9]+)_) = 1.71), sex (X²(_([0-9]+)_) = 0.86), or the interaction between day and sex (X²(_([0-9]+)_) = 0.61). For the duration of LORR, a separate analysis comparing days 1 and 4 indicated evidence of tolerance to the motor depressant effect of ethanol, as LORR duration was significantly shorter on day 4 compared to day 1 (X²(_([0-9]+)_) = 5.06; p = 0.025). No effect of sex was observed (X²(_([0-9]+)_) = 3.05). BEC levels confirmed heavy intoxication and no significant sex differences were observed (X²(_([0-9]+)_) = 1.68) (Figure 2A). For females, BEC was 305.9 ± 21.59 mg/dL on day 1 and 293.5 ± 12.49 mg/dL on day 4 (mean ± standard error), with no difference between days (X²(_([0-9]+)_) = 0.33). For males, BEC was 272.7 ± 19.14 mg/dL on day 1 and 284.77 ± 14.23 mg/dL on day 4, with no difference between days (X²(_([0-9]+)_) = 0.26). The LORR procedure for Experiments 2 and 3 yielded similar results, regarding latency and duration (Figures S1). When combining Experiments 2 and 3, the analysis of the latency to LORR showed no significant effect of day on latency to LORR in females (X²_(3)_ = 6.91) or males (X²_(3)_ = 6.89) (Figure 2B). Analysis of the LORR duration revealed a significant effect on LORR duration in both sexes (Females: X²_(3)_ = 16.52, p = 0.001; Males: X²_(3)_ = 12.53, p = 0.006), with all days differing from day 1 (Figure 2C).

**Figure 2:**
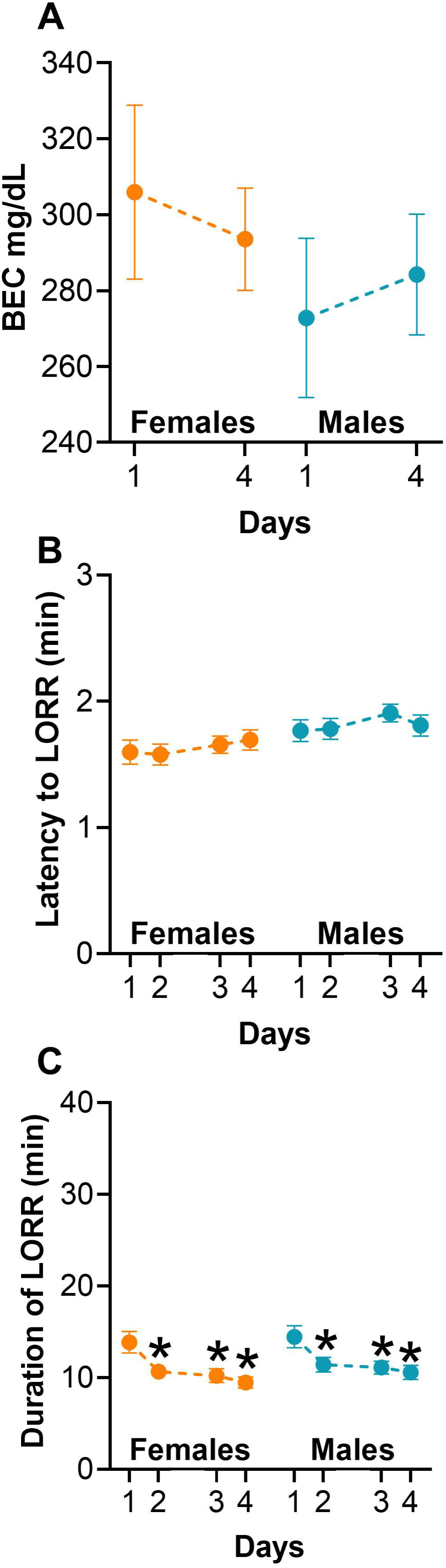
Repeated ethanol administration during adolescence induced high BEC levels and tolerance to the depressant effects of ethanol. All graphs show means ± standard error means (SEM). Orange represents data from females and blue from male mice. (*) indicates difference from day 1 (p < 0.05). **(A)** BEC level (mg/dL) at the time mice recovered the reflex in Experiment 1 (N = 7-8/sex). **(B)** Latency and **(C)** Duration of LORR in minutes for experiments 2 and 3 (N = 27-31/sex). LORR: loss of righting reflex. *: p < 0.05 compared to day 1.

### 3.2 Experiment 2: Effects of early adolescent intoxication on anxiety-like, exploratory and risk assessment behaviors responses to an ethanol challenge in adulthood

#### 3.2.1 LDB

Initially, we analyzed variables traditionally used to infer anxiety-like behaviors for the LDB test. Analysis of the latency to enter the light compartment did not show a significant effect of the factor sex (X²_(1)_ = 1.46). Separate analyses of female and male mice revealed no significant effect of the factor intoxication (adolescent saline or ethanol exposure) (Females: X²_(1)_ = 0.00; Males: X²_(1)_ = 3.80, p = 0.05), pretest (saline or ethanol administration prior to testing) (Females: X²_(1)_ = 1.31; Males: X²_(1)_ = 3.83, p = 0.05), or the interaction (Females: X²_(1)_ = 1.71; Males: X²_(1)_ = 2.66) (Figure 3A). It is worth noting that in males, there was a marginally significant effect of intoxication and pretest factors, suggesting a trend toward increased latency to enter the light side in intoxicated mice treated with saline before the test (p = 0.05). The most common variable used to indicate anxiety-like behavior is the percentage of time spent on the light side. The analysis did not detect significance for the factor sex (X²_(1)_ = 0.47). Analyses for each sex did not indicate significance of intoxication (Females: X²_(1)_ = 2.15; Males: X²_(1)_ = 0.00), pretest (Females: X²_(1)_ = 0.43; Males: X²_(1)_ = 3.70), or interaction (Females: X²_(1)_ = 0.99; Males: X²_(1)_ = 0.12) (Figure 3B). Thus, ethanol intoxication or pretest treatment do not affect anxiety-like behavior in the LDB test.

**Figure 3:**
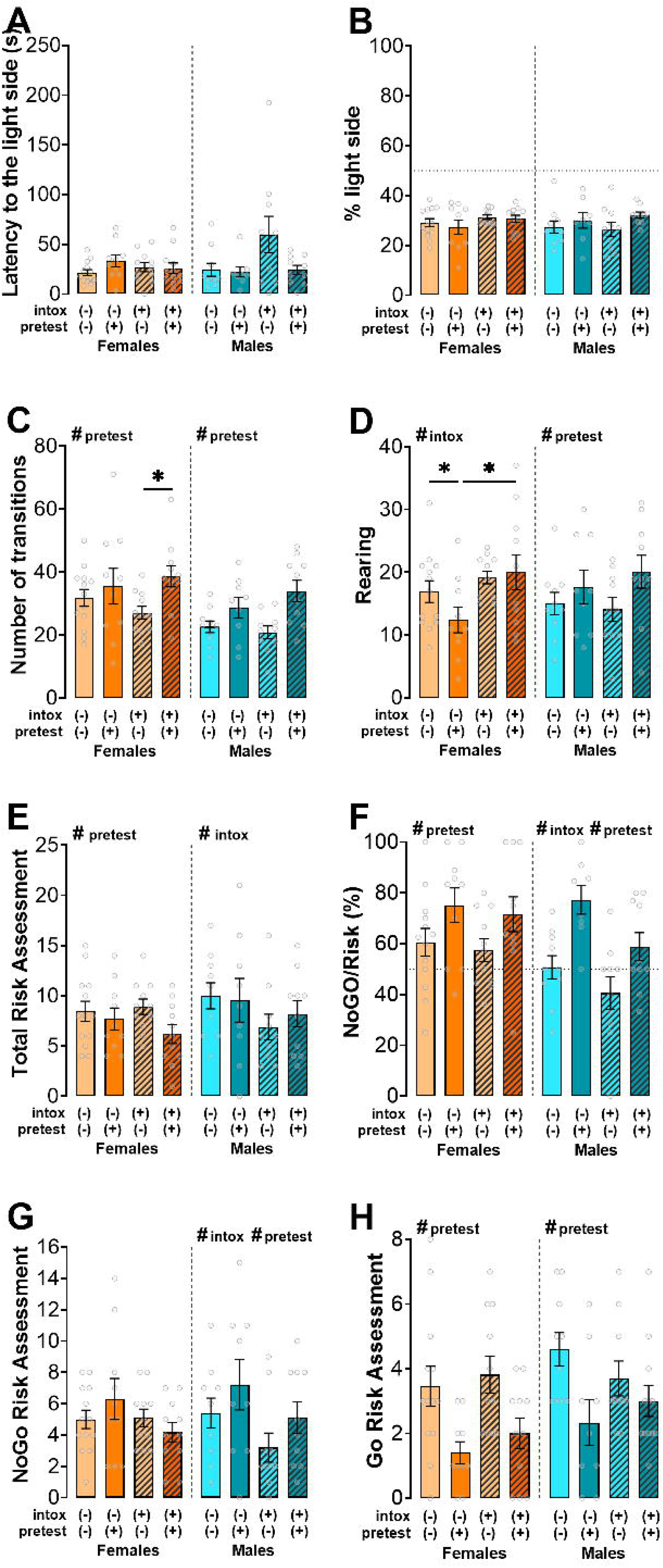
Anxiety-like, locomotion and risk assessment behaviors in the light–dark box (LDB). All graphs show means ± standard error means (SEM). The orange-toned graphs represent data from females, and the blue-toned graphs represent data from males. Hatched bars represent the intoxicated groups, which received ethanol during adolescence. Light colors (light orange, and light blue) represent animals that received saline in adulthood, while dark colors (orange, and blue) represent animals that received ethanol in adulthood. **(A)** Latency to pass to the light compartment. **(B)** Percentage of time spent in the light compartment. **(C)** Number of transitions between compartments. **(D)** Number of rearings. **(E)** Number of total risk assessments. **(F)** Percentage of NoGo risk assessments relative to the total risk assessments. **(G)** NoGo risk assessments. **(H)** Go risk assessments. N = 10-13/group/sex. intox: intoxicated during adolescence. #: p < 0.05 for main factor significance. *: p < 0.05 *post hoc* analysis.

We analyzed exploratory behaviors by counting the transitions between compartments and rearing behaviors in the light side of the LDB. The analysis of transitions detected a significant effect of sex (X²_(1)_ = 34.46, p < 0.001), with females exhibiting a higher number of transitions compared to males. Separate analyses of female data showed no effect for intoxication (X²_(1)_ = 0.38), but significant pretest (X²_(1)_ = 20.27, p < 0.001) and interaction effects (X²_(1)_ = 5.77, p = 0.016). Among intoxicated females, pretest ethanol induced more transitions than pretest saline (p < 0.001). For males, no effect was detected for intoxication (X²_(1)_ = 0.21) or interaction (X²_(1)_ = 3.49), but the factor pretest was significant (X²_(1)_ = 30.60, p < 0.001), indicating that pretest ethanol increased transitions in males independently of the intoxication during early adolescence (Figure 3C). Considering rearing counts, the analysis did not reveal a significant sex effect (X²_(1)_ = 0.10). When analyzing females, no effect of pretest was observed (X²_(1)_ = 3.33), but there were significant effects of intoxication (X²_(1)_ = 16.86, *p* < 0.001) and interaction (X²_(1)_ = 5.74, p = 0.017). In control mice, pretest ethanol decreased rearings (p = 0.034), but this effect was not observed in previously intoxicated females. The pretest ethanol effect on rearing was higher in intoxicated mice (p < 0.001). In contrast, when analyzing males, there was no effect of intoxication (X²_(1)_ = 0.18) or interaction (X²_(1)_ = 1.46), but the pretest factor was significant (X²_(1)_ = 10.79, p = 0.001). Pretest ethanol increased rearing compared to saline in males (p < 0.001) (Figure 3D). Altogether, these results suggest that pre‐test ethanol enhanced exploratory activity across both sexes, although it decreased rearing in non‐intoxicated females, with adolescent intoxication selectively modulating this response in females.

An important goal of this study was to better understand the effects of ethanol exposure during early adolescence on risky decision-making in adulthood, using risk assessment measures. The factor sex was not significant (X²_(1)_ = 1.59). Analysis of the female dataset revealed no intoxication (X²_(1)_ = 1.09) or interaction effects (X²_(1)_ = 2.30), but a pretest significant effect (X²_(1)_ = 5.73, p = 0.017). Females who received ethanol before the test performed fewer risk assessments than those treated with saline (p = 0.017). Males otherwise showed no effect of the pretest factor (X²_(1)_ = 0.35) or interaction (X²_(1)_ = 0.40), but the intoxication effect was significant (X²_(1)_ = 5.54, p = 0.019). Intoxicated adolescent male mice exhibited significantly fewer risk assessment behaviors compared to controls, regardless of pretest drug administration (p = 0.019) (Figure 3E). We also evaluated the relative percentage of NoGo behaviors (NoGo/Risk, %) within the total risk assessment (Go + NoGo). The analysis revealed no significant effect of sex (X²_(1)_ = 3.30). Female data analysis revealed no effect of the intoxication (X²_(1)_ = 0.46) or interaction (X²_(1)_ = 0.53), but a significant effect of the pretest factor (X²_(1)_ = 43.49, p < 0.001). Administration of ethanol prior to the test induced more NoGo risk assessment than the administration of saline in female mice. On the other hand, the male dataset analysis revealed significant effects of both pretest (X²_(1)_ = 67.84, p < 0.001) and adolescent intoxication (X²_(1)_ = 45.41, p < 0.001), but no interaction (X²_(1)_ = 1.71) (Figure 3F). Although pretest ethanol induced a higher proportion of NoGo responses relative to total risk assessments in females and males, intoxication induced a lower proportion of this behavior, indicating that the adolescent exposure to high doses of ethanol tends to induce male mice to riskier decision-making. For NoGo risk assessment, the analysis showed no significant effect of sex (X²_(1)_ = 0.02). Analysis of female mice data did not show significant effects of adolescent intoxication (X²_(1)_ = 2.18), pretest (X²_(1)_ = 0.00) or interaction (X²_(1)_ = 0.59). For males, there was a significant effect of intoxication (X²_(1)_ = 9.16, p = 0.002) and pretest (X²_(1)_ = 6.88, p = 0.009), but no interaction (X²_(1)_ = 0.36). Male mice that received ethanol pretest performed more NoGo risk assessments than saline-treated controls (p = 0.009). Additionally, intoxicated males exhibited fewer NoGo risk assessment compared to control (p = 0.003) (Figure 3G). We also analyzed stretch-attend postures and head-out behaviors separately (Figure S2). The analysis of NoGo stretches revealed a significant effect of sex (X²_(1)_ = 6.03, p = 0.014), with males performing significantly fewer NoGo stretches than females (p = 0.015). Among females, no significant differences were observed. Among males, the analysis detected a significant effect of intoxication (X²_(1)_ = 5.21, p = 0.022) and pretest (X²_(1)_ = 8.80, p = 0.003) (Figure S2A). Considering NoGo head-outs, the analysis revealed the sex factor as significant (X²_(1)_ = 6.29, p = 0.012), as females did more NoGo head outs than males. The analysis for each sex did not indicate any specific differences (Figure S2B). We also analyzed Go-type risk assessment behaviors, which occurred less frequently than NoGo risk assessment behaviors. The analysis showed a significant effect of sex (X²_(1)_ = 5.80, p = 0.016), with males performing more Go risk assessments than females (p = 0.017). Among females, only the pretest factor was significant (X²_(1)_ = 17.70, p < 0.001), revealing that pretest ethanol administration induced fewer Go risk assessments compared to saline (p < 0.001). Similar effects were observed in males (intoxication: X²_(1)_ = 0.02; pretest: X²_(1)_ = 5.76, p = 0.016; interaction: X²_(1)_ = 2.03) (Figure 3H). Considering the Go stretches, the analysis showed a significant effect of sex (X²_(1)_ = 10.06, p = 0.002), with males performing more Go stretches than females (p = 0.002). The analysis detected a significant effect of pretest ethanol administration in females (X²_(1)_ = 13.55, p < 0.001) and males (X²_(1)_ = 4.43, p = 0.035) (Figure S2C). Regarding the number of Go head-outs, no significant sex effect was observed. Among females, a significant pretest effect was found (X²_(1)_ = 6.67, p = 0.010), indicating that ethanol prior to the test induced fewer Go head outs when compared to saline-treated mice. No differences were found among males (Figure S2D).

We analyzed the transition probabilities to and from risk assessment behaviors,and used them to construct Markov Chain Models that illustrate the behavioral organization of each experimental group (Figure 4). In these models, the size of each circle reflects the frequency of a given behavioral state, while the color of arrows indicate the probability of transitioning from one behavior to another. Data from female and male mice were pooled. Notably, mice that received acute ethanol prior to testing displayed Go risk-assessment behaviors (Go stretch) in less than 2% of the total session time, which accounts for their absence in the visual model. However, even though this behavior was rare, acute ethanol significantly increased the likelihood of key transitions involving it: specifically, from the dark compartment to Go stretch (χ²_(1)_ = 5.87, p = 0.015) and from Go stretch to the light compartment (χ²_(1)_ = 4.55, p = 0.033). This suggests that acute ethanol exposure reshaped the *sequence* of behaviors rather than their frequency alone, promoting more direct transitions toward exploration. By contrast, no significant changes were observed in transition structure following early adolescent intoxication, indicating that only the acute ethanol challenge disrupted the organization of exploratory and defensive behavior in this task. Overall, this analysis reveals that pretest ethanol simplified and redirected the behavioral flow in the Light-Dark Box, supporting a less deliberative approach strategy.

**Figure 4.**
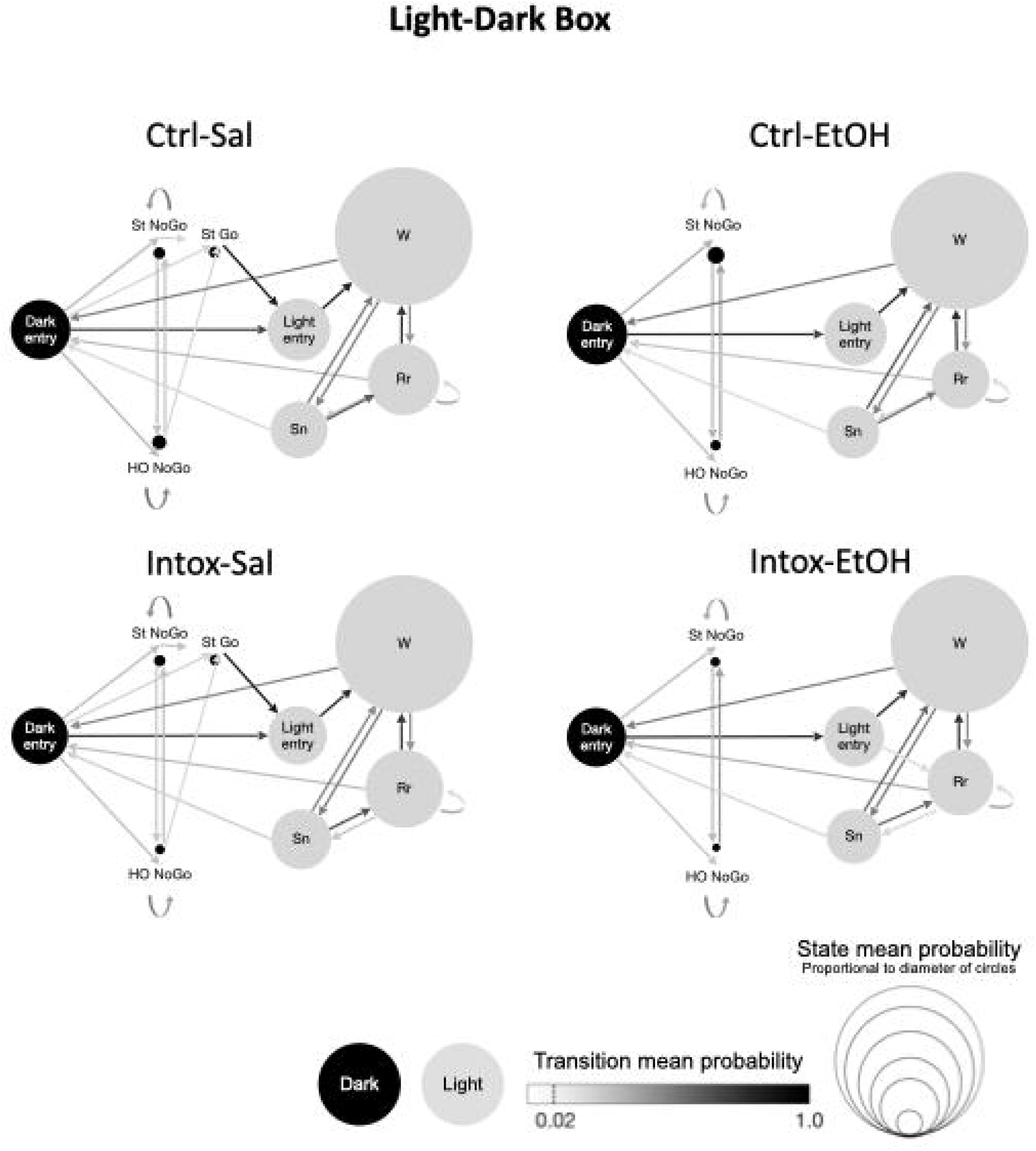
Markov Chain Models depicting how behavioral states are organized and linked during the Light–Dark Box (LDB) test. Black circles indicate behaviors performed in the dark compartment and gray circles indicate behaviors performed in the light compartment. Larger circles correspond to behaviors that occurred more frequently, while darker arrows represent transitions with higher probability. These models highlight how risk‐assessment and exploratory actions are chained together within each experimental group. Behavioral codes: HO = head‐out; St = stretch; Rr = rear; Sn = sniff; W = walk. N = 20-26/group.

#### 3.2.1 EPM

Mice were also tested in the EPM. We first analyzed variables traditionally used to infer anxiety-like behaviors. The analysis of the percentage of time spent in the open arms did not show a significant effect of the factor sex (X²_(1)_ = 0.27). For female mice, the analysis did not detect significant effects of intoxication (X²_(1)_ = 0.17) or interaction (X²_(1)_ = 0.89), but the pretest factor was significant (X²_(1)_ = 12.52, p < 0.001). Similar results were observed for males (intoxication: X²_(1)_ = 2.48; pretest: X²_(1)_ = 32.39, p < 0.001; interaction: X²_(1)_ = 2.46). Pretest ethanol produced an anxiolytic-like effect, as mice spent significantly more time in the open arms compared to those that received pretest saline (p < 0.001) (Figure 5A). We also evaluated the open arm entries and observed no significant effect of sex (X²_(1)_ = 2.55). Female results revealed a significant effect of pretest (X²_(1)_ = 10.48, p = 0.001), but no effect of intoxication (X²_(1)_ = 0.33) or interaction (X²_(1)_ = 0.45). Similar results were observed for males (intoxication: X²_(1)_ = 0.28; pretest: X²_(1)_ = 43.52, p < 0.001; interaction: X²_(1)_ = 0.49). Thus, pretest ethanol increased open arm entries when compared to saline (p < 0.001) in both sexes (Figure 5B). Overall, acute ethanol induced the expected anxiolytic effect in the EPM.

**Figure 5:**
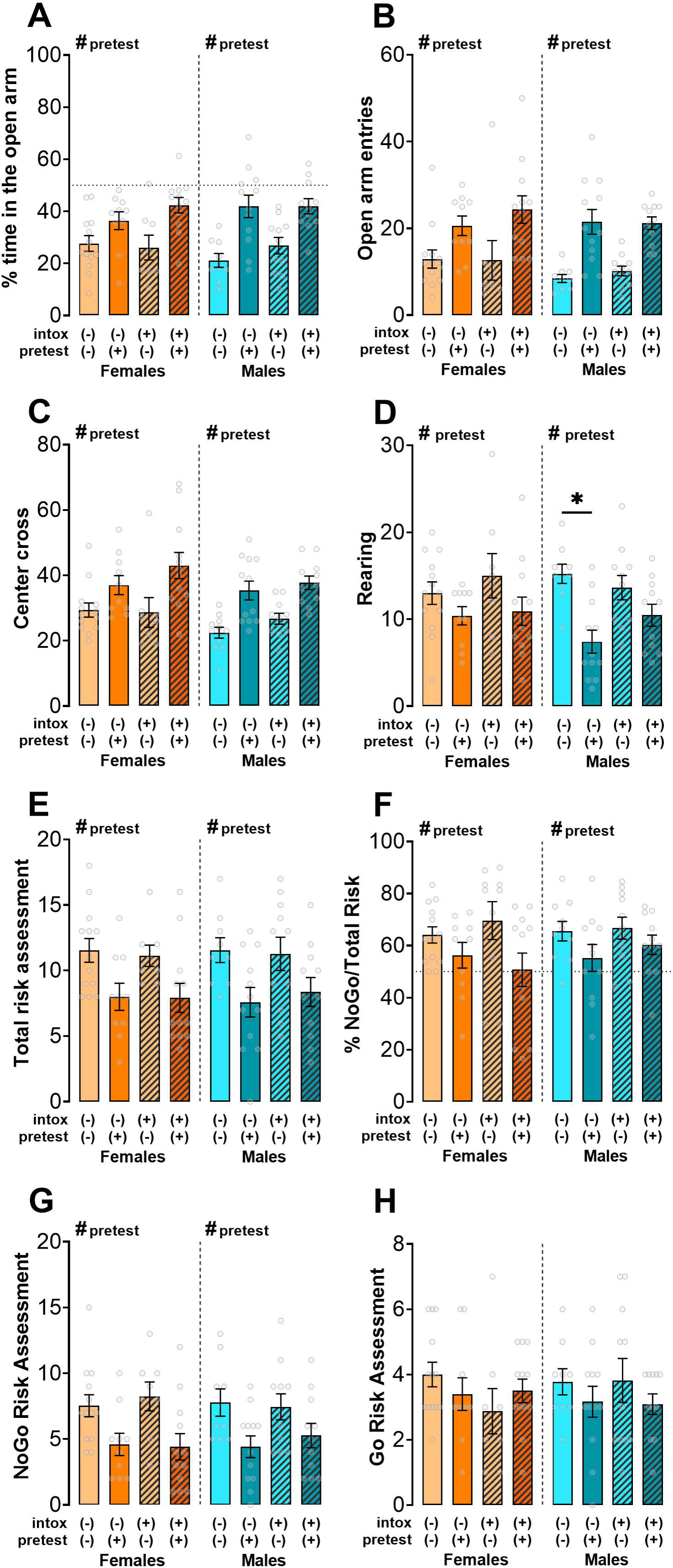
Anxiety-like, locomotion and risk assessment behaviors in the elevated plus maze (EPM). All graphs show means ± standard error means (SEM). The orange-toned graphs represent data from females, and the blue-toned graphs represent data from males. Hatched bars represent the intoxicated groups, which received ethanol during adolescence. Light colors (light orange, and light blue) represent animals that received saline in adulthood, while dark colors (orange, and blue) represent animals that received ethanol in adulthood. **(A)** Percentage of time spent in the open arms. **(B)** Number of entrances in the open arms. **(C)** Number of center crosses. **(D)** Number of rearings. **(E)** Number of total risk assessments. **(F)** Percentage of NoGo risk assessments relative to the total risk assessments. **(G)** NoGo risk assessments. **(H)** Go risk assessments. N = 10-13/group/sex. #: p < 0.05 for main factor significance. *: p < 0.05 *post hoc* analysis.

Exploratory behavior can be inferred by the number of center crossings and rearing. The analysis of center crossings detected a sex effect (X²_(1)_ = 9.33, p = 0.002), as female mice made more center crossings than males (p = 0.002). Female data revealed no intoxication (X²_(1)_ = 1.32) or interaction (X²_(1)_ = 2.70) effects, but it showed a significant pretest effect (X²_(1)_ = 35.60, p < 0.001). For males, although no significant interaction was detected (X²_(1)_ = 0.97), intoxication (X²_(1)_ = 4.71, p = 0.030) and pretest factors (X²_(1)_ = 53.35, p < 0.001) were significant. Pretest ethanol increased center crossings compared to saline-treated mice from both sexes (p < 0.001) (Figure 5C). Considering rearing behavior, the analysis revealed no significant effect for the sex (X²_(1)_ = 1.38). When analyzing females data, no effect was observed for intoxication (X²_(1)_ = 1.17) or interaction (X²_(1)_ = 0.28), but the pretest factor was significant (X²_(1)_ = 9.34, p = 0.002), indicating that ethanol prior to the test decreased rearings compared to saline (p < 0.001). Among males, the intoxication factor was not significant (X²_(1)_ = 3.17), but the pretest factor (X²_(1)_ = 24.71, p < 0.001) and the interaction (X²_(1)_ = 4.00, p = 0.046) were significant. Pretest ethanol decreased the number of rearings only in naive mice (p < 0.001) (Figure 5D). Overall, pretest ethanol enhanced center crossings but reduced the number of rearings across the entire apparatus, an effect not observed in previously intoxicated male mice.

We also analyzed the risk assessment behaviors in the EPM. For the total amount of risk assessment, the analysis indicated no significant effect of sex (X²_(1)_ = 0.04) (Figure 4J). For females, no effect of intoxication (X²_(1)_ = 0.05) or interaction (X²_(1)_ = 0.02), but a significant pretest was observed (X²_(1)_ = 12.29, p < 0.001). Males showed similar results (intoxication: X²_(1)_ = 0.25; pretest: X²_(1)_ = 12.87, p < 0.001; interaction: X²_(1)_ = 0.25). Female and male mice that received ethanol immediately before the test performed fewer risk-assessment behaviors than controls (p < 0.001) (Figure 5E). When analyzing the percentage of NoGo risk assessments relative to the total risk assessments (% NoGo/Risk), the analyses revealed no significant effects of the factor sex (X²_(1)_ = 0.35). Among females, no effect of the factor intoxication (X²_(1)_ = 0.02) or the interaction (X²_(1)_ = 1.52), but the analysis showed a significant pretest effect (X²_(1)_ = 6.68, p = 0.01). Again, male mice showed similar results (intoxication: X²_(1)_ = 0.19; pretest: X²_(1)_ = 4.95, p = 0.026; interaction: X²_(1)_ = 0.64). Pretest ethanol induced a lower percentage of NoGo risk assessments than saline in both sexes (p < 0.05) (Figure 5F). Acute ethanol administration prior to EPM testing consistently reduced risk-assessment behaviors, particularly NoGo responses, in both sexes, regardless of prior adolescent intoxication history. Considering only the NoGo risk assessment behaviors, the analysis showed no significant effect of the factor sex (X²_(1)_ = 0.14). Among females, intoxication (X²_(1)_ = 0.04) and interaction (X²_(1)_ = 0.26) were not significant, but the pretest factor was significant (X²_(1)_ = 19.54, p < 0.001), revealing that pretest ethanol administration induced fewer NoGo risk assessments compared to pretest saline (p < 0.001). Similar effects were observed in males (intoxication: X²_(1)_ = 0.28; pretest: X²_(1)_ = 14.37, p < 0.001; interaction: X²_(1)_ = 0.84) (Figure 5G). The analysis of NoGo stretches revealed no effect of sex. Among females, the analysis revealed significant effects of pretest (X²_(1)_ = 9.82, p = 0.002), but among males no significant differences were observed (Figure S3A). Considering the NoGo head outs, the analysis showed a significant effect of sex (X²_(1)_ = 8.35, p = 0.004), as males performed more NoGo head outs than females (p = 0.004). Analyzing each sex separately, similar effects were observed for the pretest factor in females (X²_(1)_ = 9.00, p = 0.003) and males (X²_(1)_ = 13.95, p < 0.001) (Figure S3B). We also analyzed Go-type risk assessment behaviors. In this case, the analysis revealed no sex effect, and the analysis for each sex did not indicate any specific differences (Figure 5H). When analyzing Go stretches, no significant effects of sex were observed and the analysis for each sex did not indicate any specific differences (Figure S3C). The same lack of significance was observed when analyzing Go head outs for both sexes (Figure S3D).

In the EPM, we also analyzed the transition probabilities involving risk assessment behaviors and constructed Markov Chain Models to characterize the organization of behavioral sequences across groups (Figure 6). In these models, circle size represents the frequency of each behavior, while arrow color indicates the probability of transitions between behaviors. Data from female and male mice were pooled. In animals that received pretest ethanol, the models revealed a significant disruption in risk-assessment organization. Specifically, NoGo behaviors, such as head-out and stretch postures, were weakly represented or nearly absent, reflecting their low frequency (< 2%). The first-order transition probability analysis revealed robust effects of pretest ethanol exposure, with no significant effects of early adolescent intoxication. Specifically, pretest ethanol significantly reduced transitions from NoGo head out or NoGo stretch to closed-arm walking (χ²_(1)_ = 23.16, p < 0.001 and χ²_(1)_ = 12.22, p < 0.001, respectively), as well as transitions from NoGo head outs to NoGo stretch (χ²_(1)_ = 10.78, p = 0.001). Furthermore, there was a reduced persistence in NoGo stretch behavior, as shown by decreased self-transitions (χ²_(1)_ = 4.11, p = 0.043), and fewer transitions from closed-arm walking to either NoGo head-out or NoGo stretch (χ²_(1)_ = 22.78, p < 0.001 and χ²_(1)_ = 9.96, p = 0.002, respectively). Thus, in contrast to saline-treated mice, which exhibited rich and interconnected NoGo transition sequences, ethanol-pretreated mice displayed a simplified behavioral structure dominated by direct transitions between exploration and non-risk behaviors. Together, these findings indicate that acute ethanol administration markedly reduces not only the expression but also the structural integration of risk-assessment behaviors within the EPM, suggesting a breakdown in the complexity of defensive decision-making. This effect was not affected by previous early adolescent ethanol intoxications.

**Figure 6.**
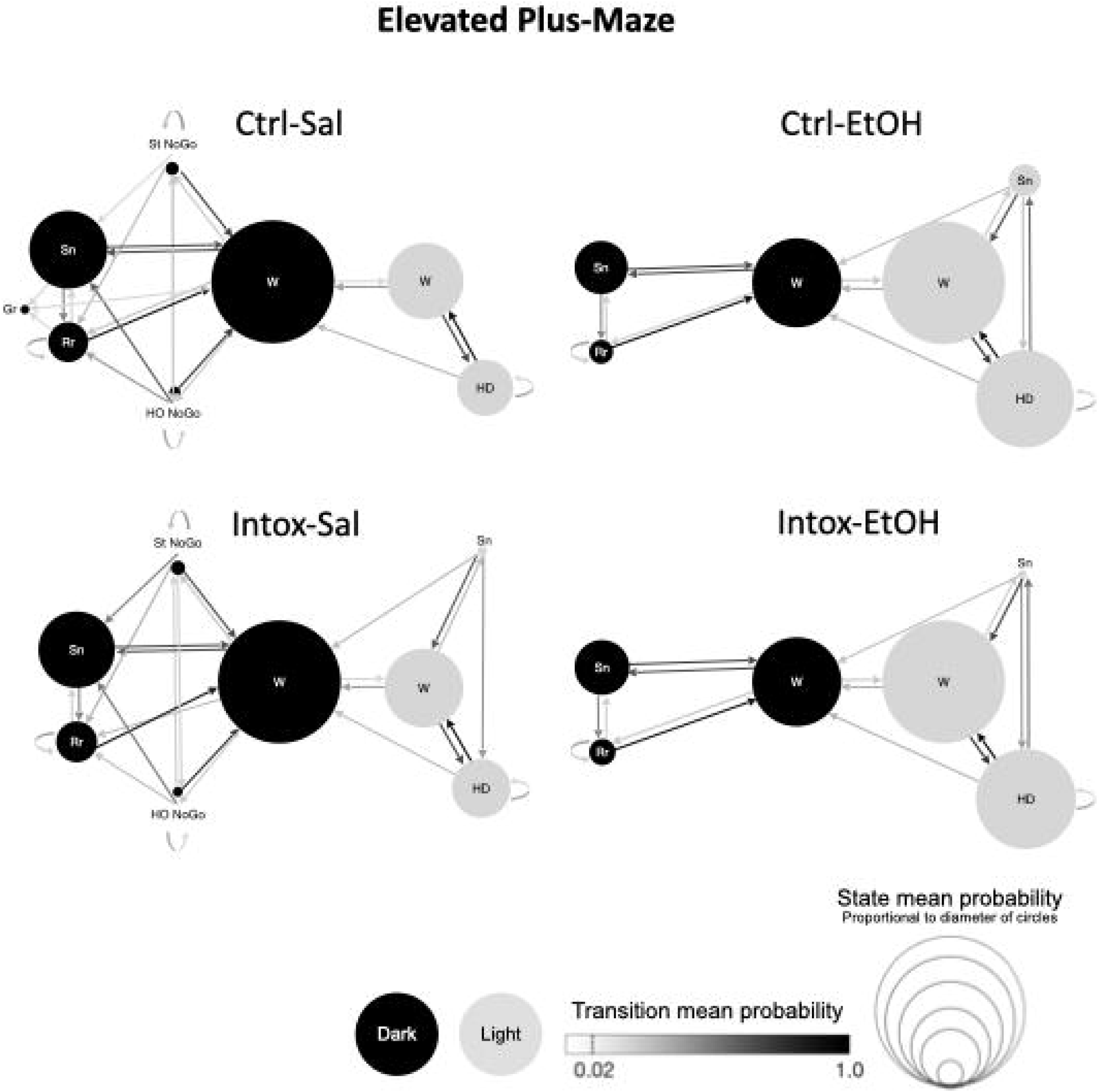
Markov Chain Models illustrating the organization and sequencing of behavioral states during the Elevated Plus Maze (EPM) test. Black circles represent behaviors occurring in the closed arms, while gray circles indicate behaviors occurring in the open arms. Circle size reflects the relative frequency of each behavioral state, with larger circles indicating more frequently expressed behaviors. Arrow color intensity represents the probability of transitioning from one behavior to another, highlighting how exploratory and risk‐assessment behaviors are chained during maze exploration. Differences in circle size and connectivity illustrate changes in the recurrence and structure of behavioral sequences across experimental conditions. Behavioral codes: Gr = grooming; HO = head‐out; St = stretch; Rr = rear; Sn = sniff; W = walk; HD = head dips. N = 20–26 per group.

**Figure 7:**
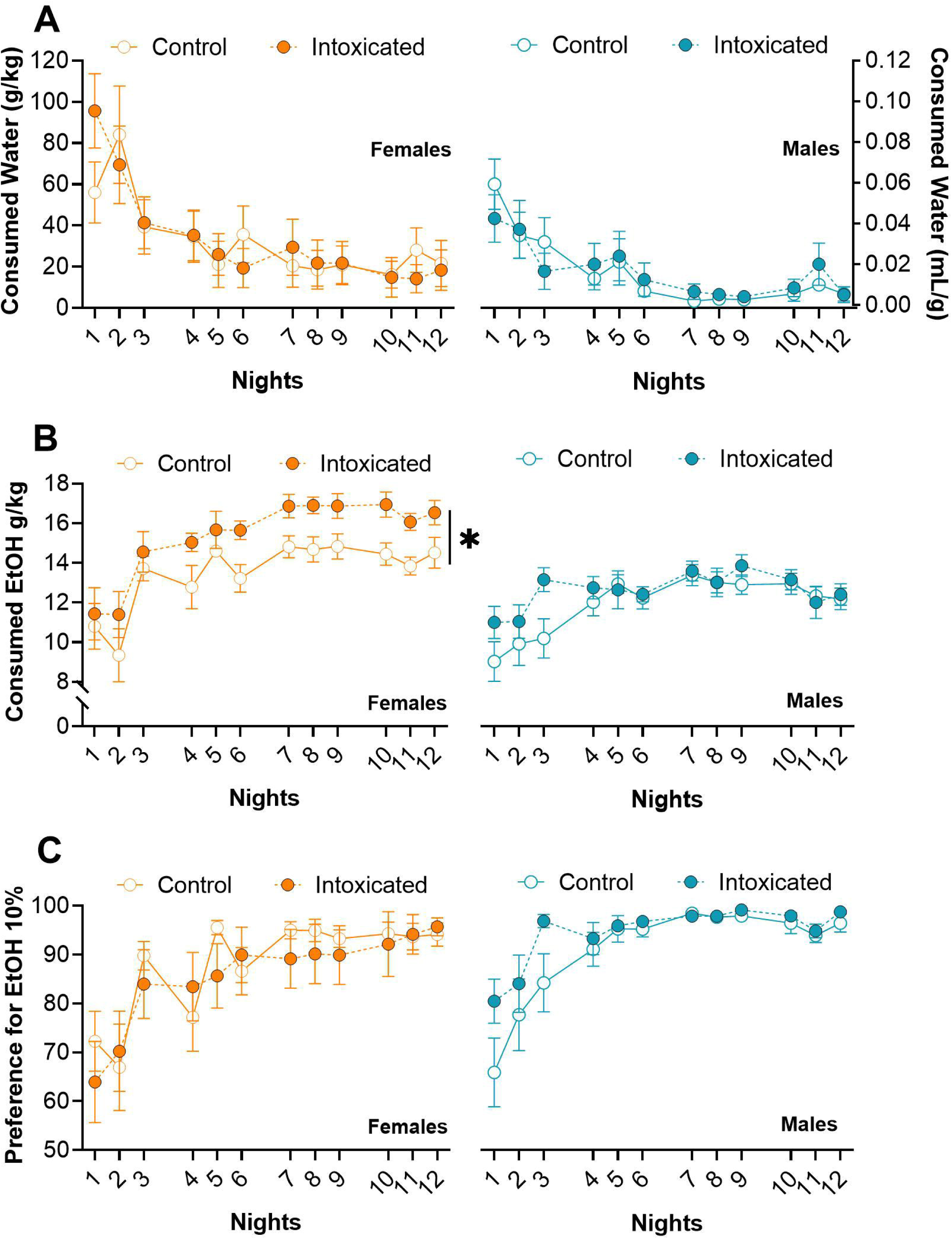
Effects of heavy intoxications during adolescence increases adulthood ethanol drinking in females. All graphs show means ± standard error means (SEM). Orange represents data from females and blue from males. White circles represent control groups, while circles filled with orange or blue represent adolescence-intoxicated groups. **(A)** Water intake (g/kg). **(B)** Ethanol intake (g/kg) from 10% (v/v) solution. **(C)** Ethanol preference. N = 17-18/group/sex. *: p < 0.05 for group factor significance.

Together, these findings indicate that pretest ethanol not only reduces the overall expression of NoGo risk assessment behaviors, as shown by the frequentist analyses, but also markedly disrupts the structure and recurrence of these behaviors in the EPM, leading to a globally reduced engagement in ethologically relevant defensive behaviors.

### 3.3 Experiment 3: Early adolescence intoxication increases adulthood voluntary ethanol consumption in females

We measured the food ingested by each mouse while they were in the IOD protocol and no difference in food intake was observed between controls and intoxicated mice (Figure S4). For water consumption, there was a significant effect of sex (X²_(1)_ = 10.09, p = 0.002), as females drank more water than males (p = 0.002) (Figure 6B). For females, no effect of group (X²_(1)_ = 1.21) or interaction (X²_(11)_ = 18.59) was observed, but the factor night was significant (X²_(11)_ = 86.39, p < 0.001), with females drinking less water on night 3 onward compared to night 1 (p < 0.001). For males, there was no effect of group (X²_(1)_ = 1.31) or interaction (X²_(11)_ = 15.19), but the factor night was significant (X²_(11)_ = 87.45, p < 0.001); males drank less water on nights 4, 6-10 and 12 compared to night 1 (p < 0.001). Overall, water intake decreased over nights in both sexes, but females consistently drank more than males.

Regarding ethanol consumption, there was a significant effect of the factor sex (X²_(1)_ = 42.88, p < 0.001), as females consumed more ethanol than males (p < 0.001) (Figure 6D). Analysis of the female dataset revealed a significant effect of night (X²_(11)_ = 92.86, p < 0.001), indicating an escalation in ethanol intake, and a significant group effect (X²_(1)_ = 21.31, p < 0.001), with previously intoxicated females consuming more ethanol than controls (p < 0.001). No significant interaction was found (X²_(11)_ = 7.46). In contrast, among males, the effect of night was significant (X²_(11)_ = 83.90, p < 0.001), indicating increased ethanol consumption over time; however, there were no significant effects of group (X²_(1)_ = 1.77) or interaction (X²_(11)_ = 16.28). Ethanol intake escalated over time in both sexes, with previously intoxicated females consuming more than controls.

Regarding ethanol preference, there was no effect of sex (X²_(1)_ = 1.78). For females, there was no group effect (X²_(1)_ = 2.88), but a significant effect of night (X²_(11)_ = 111.42, p < 0.001), with ethanol preference increasing from night 3 onward compared to night 1 (p < 0.001); and no interaction was found (X²_(11)_ = 16.41). Likewise, for males, there was no group effect (X²_(1)_ = 3.22), but a significant effect of night (X²_(11)_ = 70.76, p < 0.001), with ethanol preference emerging from night 3 onward (p < 0.001) and the interaction was not significant (X²_(11)_ = 11.40). Ethanol preference increased independent of prior intoxication history.

## 4. Discussion

Ethanol exposure can influence behavior in complex ways, depending on factors such as dose, age and sex. Therefore, it is important to investigate if and how behavioral changes take place when subjects have been previously exposed to ethanol, particularly during adolescence, and/or are under its acute effects later in life. In the present study, ethanol intoxication in early adolescence reduced NoGo risk assessment behaviors later in life in the LDB, specifically in males, while no effect was observed in the EPM. The well-established immediate anxiolytic effect of a low ethanol dose was accompanied by increased exploratory behavior and reduced male NoGo risk assessment behaviors in the EPM. Interestingly, the acute anxiolytic effect of ethanol was not observed in the LDB. However, ethanol administered prior to the LDB increased exploratory behavior in both sexes, which was accompanied by an increase in male NoGo risk assessment and a decrease in Go risk assessment behaviors in both sexes. Finally, we observed that early adolescent intoxication increased voluntary ethanol intake in adult females, an effect not observed in males.

To induce intoxication, we used a model of repeated high-dose ethanol administrations, which can lead to LORR (31,32) and may represent repeated episodes of alcohol-induced passing out in humans. Importantly, losing the righting reflex is different from sleep, as 4.0 g/kg of ethanol causes a brain activity state different from sleep in mice (33). Although LORR paradigms are well established in adult mice, relatively few studies have implemented adolescent high‐dose ethanol exposures capable of producing LORR‐level intoxication, and even fewer have explicitly assessed LORR during this developmental period in mice. We observed a shorter LORR duration in adolescents (postnatal days 36 to 45) compared to what is typically reported in the literature, even in studies using slightly higher doses of ethanol (34–36). Additionally, some mice did not lose the reflex but still presented clear behavioral intoxication. It is important to note that animals can develop LORR tolerance with repeated ethanol administrations, as indicated by faster reflex recovery. Our intoxication protocol may have induced a metabolic tolerance, as the BEC levels on days 1 and 4 were similar, even though animals showed a shorter recovery time on day 4 compared to the first day of LORR. In general, functional tolerance is observed in adult animals while metabolic tolerance is observed in adolescents (37,38). Even fewer studies have used LORR to investigate ethanol tolerance in female and male mice (36). In our study, we did not observe sex differences in the time to lose or recover the righting reflex.

Adolescent ethanol exposure has been linked to increased anxiety-like behavior, impulsivity, and risk-taking, though these effects vary depending on the exposure protocol and the method of ethanol administration (for review, see 39). The specific long-term behavioral consequences of repeated alcohol intoxications remain incompletely understood, particularly regarding sex differences and the use of structural ethoexperimental analyses to assess adult behavior in models that evoke conflict. In traditional anxiety tests, measures such as latency to enter an aversive area, percentage of time spent in that environment, and number of entries in the aversive space are commonly used as indicators of anxiety-like behavior. However, these variables are affected by the animal’s exploratory drive, as seen in changes in transitions between compartments and rearing behavior. Risk assessment, in particular, occupies a distinct but related domain. As defined by Blanchard and Meyza (2019) (40), risk assessment involves an approach–avoidance conflict in response to an ambiguous or threatening stimulus. The behavioral repertoire associated with risk assessment includes cautious engagement with the environment - such as stretch-attend postures and head outs towards the aversive environment - reflecting a balance between defensive and exploratory trends. In the present study, anxiety-like, exploratory, and risk assessment behaviors were evaluated using two behavioral tests, each involving a conflict scenario: the choice between exploring a brightly lit, novel area or remaining in a dark, safe zone (LDB), and the exposure to elevated open arms versus enclosed areas (EPM). In addition to the variables listed above, these paradigms enabled us to assess how previous ethanol exposure affected decision-making under potential threat, as we propose two categories of risk assessment behaviors: those followed by a return to the safe environment (NoGo) and those followed by taking the risk to enter the aversive environment (Go), allowing us to study risk-based decision-making.

The LDB test is traditionally used to evaluate escaping behavior, by measuring the latency of rodents to pass from the light side to the dark side of the box (41). Using this protocol, Younis and colleagues reported that male and female mice exposed to voluntary ethanol drinking during adolescence did not differ in the time to escape the light side, although mice ran faster to the dark side after an ethanol i.p. administration immediately before the test (42). In rats, only female adolescents exposed to ethanol showed decreased latency to enter the dark side when they were adults (43). We used an adapted version of the LDB: an “avoid” paradigm rather than an “escape” one, which consisted of allocating the animal first to the dark side of the apparatus, allowing them to evaluate the risk of going to the light side (40). Intoxication and pretest ethanol administration did not change latency to enter the light compartment or percentage of time spent in the light side of the LDB. Previous studies have reported that adolescent ethanol exposure increases avoidance of the aversive (light) compartment, especially in male rats (45–47). Interestingly, female mice that have been intoxicated during adolescence show a stimulant response after acute ethanol, exhibiting a greater number of transitions than non-intoxicated mice. The present study also shows that male mice intoxicated during adolescence exhibited fewer NoGo risk assessment behaviors in adulthood, suggesting a greater propensity for risk-taking. Regarding acute ethanol administration prior to testing, ethanol increased the relative percentage of NoGo risk assessment behaviors, primarily due to a reduction in Go risk assessment behaviors in both sexes. We also did not observe substantial effects on the overall behavioral structure measured in the LDB; however, this conclusion should be interpreted with caution, as behaviors in the dark compartment were not evaluated due to methodological constraints.

The EPM is a widely used test for assessing anxiety-like behavior in rodents (48). In this test, increases in anxiety-like behavior in adulthood following adolescent ethanol exposure have been reported in both rats (39) and mice (49,50) (review in 22). However, our results suggest that in C57BL/6 mice, four ethanol intoxications during early adolescence did not alter anxiety-like, exploratory, or risk assessment behaviors in adulthood, regardless of sex. One important point to consider is that mice were tested in the LDB first, which could have influenced their behavior in the EPM. Nonetheless, similar to our results, Younis et al. (42) reported that voluntary ethanol drinking during adolescence did not affect the latency to enter or time spent in the open arms of the EPM in adulthood. Regarding the effects of a pretest low dose of ethanol, we found an increased time spent in the open arms, indicative of the well established (42,51) anxiolytic acute effect of ethanol. Interestingly, the acute anxiolytic effect of ethanol is followed by increases in exploration and decreases in NoGo risk assessment behaviors. These effects profoundly change the behavior structure, as displayed on the Markov Model.

The Markov chain analysis adds an important layer of interpretation to the results by allowing us to examine the structure of behavior rather than only each behavior. Unlike traditional ethological measures, which treat behaviors as isolated, discrete events, Markov models quantify how animals transition from one behavioral state to another, revealing how manipulations alter the organization, recurrence, and sequencing of defensive and exploratory strategies. When using this approach in tests like the LDB and the EPM, it captures the dynamic nature of the approach-avoidance conflict: behaviors do not occur randomly, but follow predictable probabilistic paths that reflect the animal’s ongoing evaluation of threat, safety, and uncertainty. In this sense, Markov modeling moves beyond classical metrics to reveal deeper organizational changes in defensive behavior that would otherwise remain undetected.

The Markov models constructed for both the LDB and EPM tests offer compelling evidence that acute ethanol exposure prior to testing disrupts not only the amount but also the structure of risk assessment behavior. Standard analyses showed that ethanol reduced the total duration of these behaviors, but the Markov analysis revealed a deeper layer: ethanol profoundly altered the transition architecture of behavioral sequences. In the LDB, pretest ethanol specifically reshaped transitions surrounding Go-type risk assessment behaviors, changing how these events interfaced with exploratory patterns. In the EPM, ethanol exposure led to a stark reduction in the frequency and interconnectivity of NoGo behaviors, fragmenting what is normally a cohesive and nuanced defensive strategy. The result was a simplified behavioral sequence marked by more direct transitions between exploration and passive states. By showing that ethanol reduced the recurrence and flexibility of risk-assessment transitions, these models suggest that acute ethanol simplifies or destabilizes the behavioral system, producing a less flexible and less adaptive defense. This degradation in behavioral structure may reflect an impaired ability to navigate conflicting motivational cues, such as threat versus exploration, and resonates with findings in humans where alcohol acutely impairs complex decision-making under uncertainty. Importantly, ethanol exposure during early adolescence did not produce this level of behavioral reorganization in adulthood, suggesting that acute effects, rather than long-term developmental changes, may be the primary drivers of disrupted sequencing. Overall, these results demonstrate how Markov models can uncover subtle yet functionally meaningful reorganizations in behavioral flow, offering new insights into how ethanol compromises adaptive choice-making at the structural level of action selection.

It is noteworthy that both Younis et al (42) and the present study involved intraperitoneal injection administered prior to LDB and/or EPM testing. Even saline, used as a control, is a procedural manipulation that can influence behavioral outcomes compared to experiments in which there is no pretest injection (52). Additionally, one limitation of this study is the lack of BEC data at the behavioural challenges timepoint, as the adolescence exposure could affect BEC to 1,2 g/kg ethanol later in life. Future studies could investigate the long-term behavioral effects of adolescent ethanol exposure in the absence of pretest injections, as well as potential metabolic adaptations resulting from such early-life intoxication.

Repeated ethanol-induced intoxications during adolescence, along with the development of tolerance to ethanol’s sedative effects, have been shown to influence drinking behavior later in life (4,46,53). While many adolescent ethanol exposure protocols have been shown to increase ethanol intake in adulthood (44,54), this outcome is not consistently replicated across studies (55,56). In our study, intoxicated female mice consumed significantly more ethanol than non-intoxicated females, suggesting a sex-specific vulnerability to risky drinking patterns. It is broadly replicated that female mice drink more ethanol than males, which already points to possibly a more prone development of increased drinking. Our results further highlight the importance of considering both sex and developmental timing when investigating the long-term effects of adolescent ethanol exposure. Literature from both rodent and human studies supports robust sex differences in behaviors related to addiction (57,58). Moreover, the specific timing of ethanol exposure - whether during early, mid, or late adolescence - can result in distinct behavioral outcomes later in life (53,59).

The mechanisms by which early alcohol use increases risky decision-making, including alcohol abuse, are yet to be fully clarified. Among youth, drinking to the point of blackout or even passing out is often viewed as socially acceptable or even desirable - used as a form of celebration or a coping mechanism for stress (60–63). It is therefore essential to characterize how high-intensity ethanol exposure during adolescence interferes with the drug’s effects in adulthood, especially considering that adolescents differ neurobiologically in their sensitivity to ethanol’s sedative actions due to developmental variations, such as in inhibitory neurotransmission and beyond. Our findings suggest that heavy intoxication during early adolescence can interact with adulthood ethanol exposure to change risk assessment behaviors in males, and enhance ethanol consumption in females. Future studies should expand on our findings by comparing different developmental windows of ethanol exposure (i.e., early, mid, or late adolescence) to investigate how timing interacts with sex to shape long-term behavioral outcomes, with particular attention to risky decision-making, and also assess how these sex- and age-dependent effects may translate into heightened vulnerability to risky drinking trajectories and alcohol use disorders in humans.

## 5. CRediT authorship contribution statement

Leticia Souza Pichinin: Conceptualization, Data curation, Formal analysis, Investigation, Methodology, Visualization, Writing – original draft, Writing – review and editing. Myrna Guernelli: Data curation, Formal analysis, Visualization, Writing – review and editing. Marianna Nogueira Cecyn: Conceptualization, Data curation, Formal analysis, Visualization, Software, Writing – review and editing. Beatriz Deo Sorigotto: Conceptualization, Data curation, Formal analysis, Visualization, Writing – review and editing. Karina Possa Abrahao: Conceptualization, Data curation, Formal analysis, Funding acquisition, Methodology, Project administration, Resources, Supervision, Validation, Writing – original draft, Writing – review and editing.

## 8. Acknowledgments

We kindly thank Dr José Luiz da Costa and Aline Martins, from CIATox Unicamp for their collaboration to measure Blood Ethanol Concentration (BEC) analyses. We also thank Paula Mendonça Camargo Eduardo and Dr Mariana Cardoso Melo for their assistance with the IOD experiment.

## 9. Data availability

Data will be made available upon request.

## 10. Declaration of generative AI and AI-assisted technologies in the writing process

During the preparation of this work the author(s) used ChatGPT in order to assist with English grammar only. After using this tool/service, the authors reviewed and edited the content as needed and take full responsibility for the content of the publication.

