## Supplemental Material for "Repeated Early Adolescence Ethanol Intoxication Promotes Riskier Decision-Making in Adult Males and Increases Drinking in Adult Female mice"

**Table 1: List of analyzed behaviors during the LDB and EPM tests.**

| **BEHAVIOR** | **CATEGORY** | **DEFINITION** |
| --- | --- | --- |
| REARING | Exploratory | The mouse assumes a vertical position, supporting itself on its hind legs while the front legs either move in the air or lean against a wall. |
| HEAD DIP | Exploratory | Downward visual screening that happens when the mouse is in one of the elevated open arms of the plus maze. |
| WALK | Exploratory | The mouse moves around and changes position. Includes non-specific body movements such as turning or running across the apparatus. |
| IMMOBILE SNIFF | Exploratory | Rhythmic inhalation and exhalation of air through the nose while the mouse stands still. |
| GROOMING* | Self-care | The mouse rubs any part of the face with circular movements of the forepaws, licks its body and/or its forepaws and back paws. |
| LIGHT SIDE ENTRY | Anxiety | The mouse enters the light side of the LDB with its four paws. |
| DARK SIDE ENTRY | Anxiety | The mouse enters the dark side of the LDB with its four paws. |
| OPEN ARM ENTRY | Anxiety | The mouse moves from the central platform to an open arm. |
| CLOSED ARM ENTRY | Anxiety | The mouse moves from the central platform to a closed arm. |
| HEAD OUT^#^ | Risk assessment | While in a protected area (dark side or closed arms), the mouse extends only its head towards the aversive area (brighter/light side or open arms), then returns to the original position (NoGo) or enters the aversive area (Go). |
| STRETCH^@^ | Risk assessment | Forward extension of the head and one or both forepaws from a protected area to an aversive area, followed by retraction to the original position (NoGo) or enters the aversive area (Go) |

* Occurred less than 1% of the time during the LDB and EPM tests, and therefore was not evaluated/analyzed.

^#^ In the EPM, head out and stretch postures occur in the closed arms or central platform towards the open arm.

^@^ In the LDB, head out and stretch behaviors occur when the animal is on the dark side of the box and extends its body towards the light side.

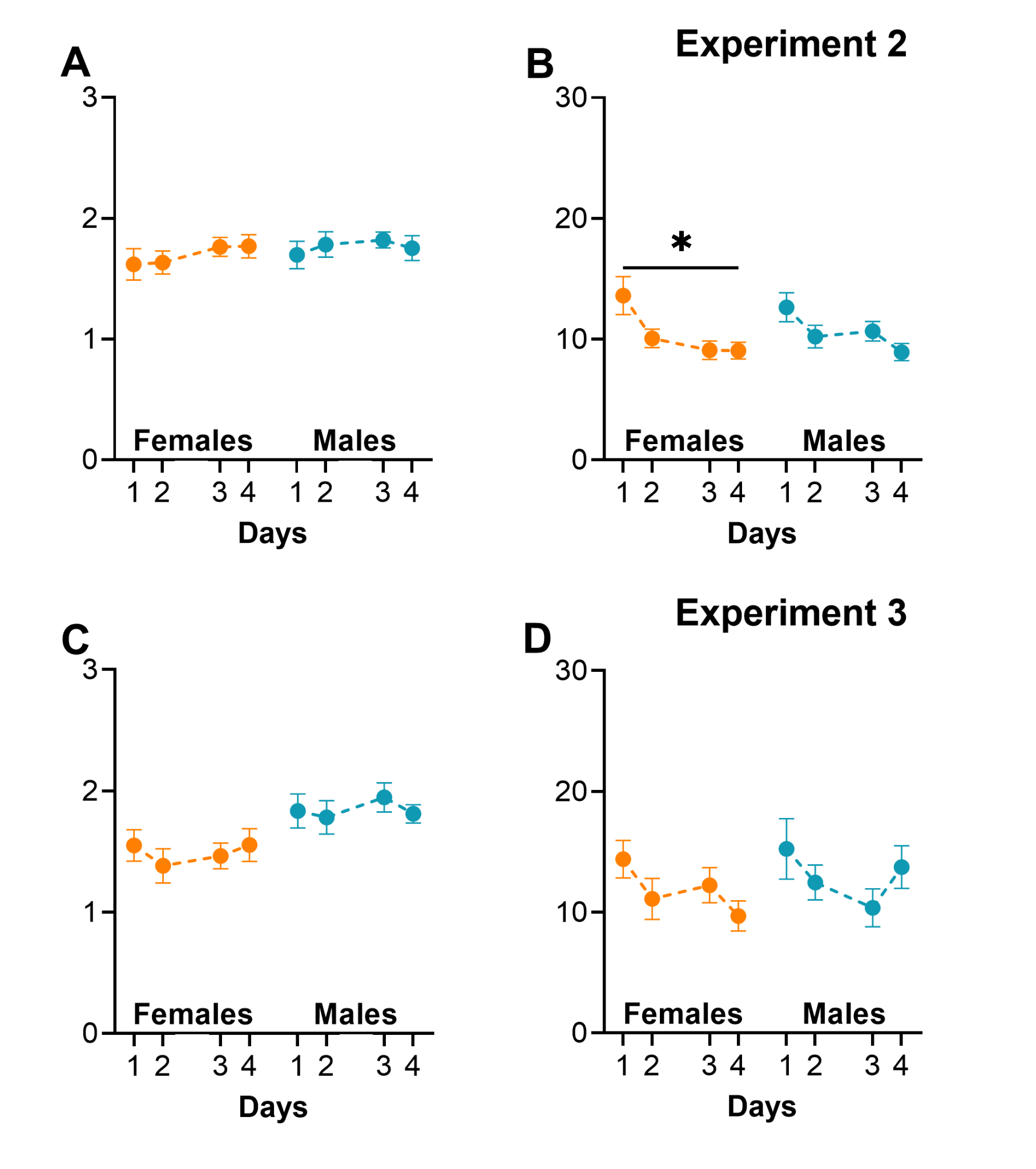

**Figure S1: Latency and Duration data of LORR from Experiment 2 and 3.** All graphs show means ± standard error means (SEM). Orange represents data from females and blue from males. (*) indicates a difference from day 1 (p < 0.05). **(A)** For latency to LORR, there was an effect of day (X²_(3)_ = 10.27, p < 0.016) but no specific difference was detected by the post hoc analysis. Additionally, there was no effect of sex. **(B)** The analysis of the duration of LORR showed a significant effect of day (X²_(3)_ = 14.26, p < 0.003) with a shorter duration on days 3 and 4 compared to day 1, indicating the development of tolerance. No significant effect of sex was observed. **(C)** Analysis of the latency to lose reflex indicated no significant effect of day or sex. **(D)** The analysis of the duration until recovery of the reflex indicated a significant effect of day (X²_(3)_ = 7.98, p < 0.047), with a shorter duration on day 4 compared to day 1. No significant effect of sex was observed. LORR: loss of righting reflex. Experiment 2: n = 10-13/group/sex. Experiment 3: n = 17-18/group/sex. *: p < 0.05 compared to Day 1.

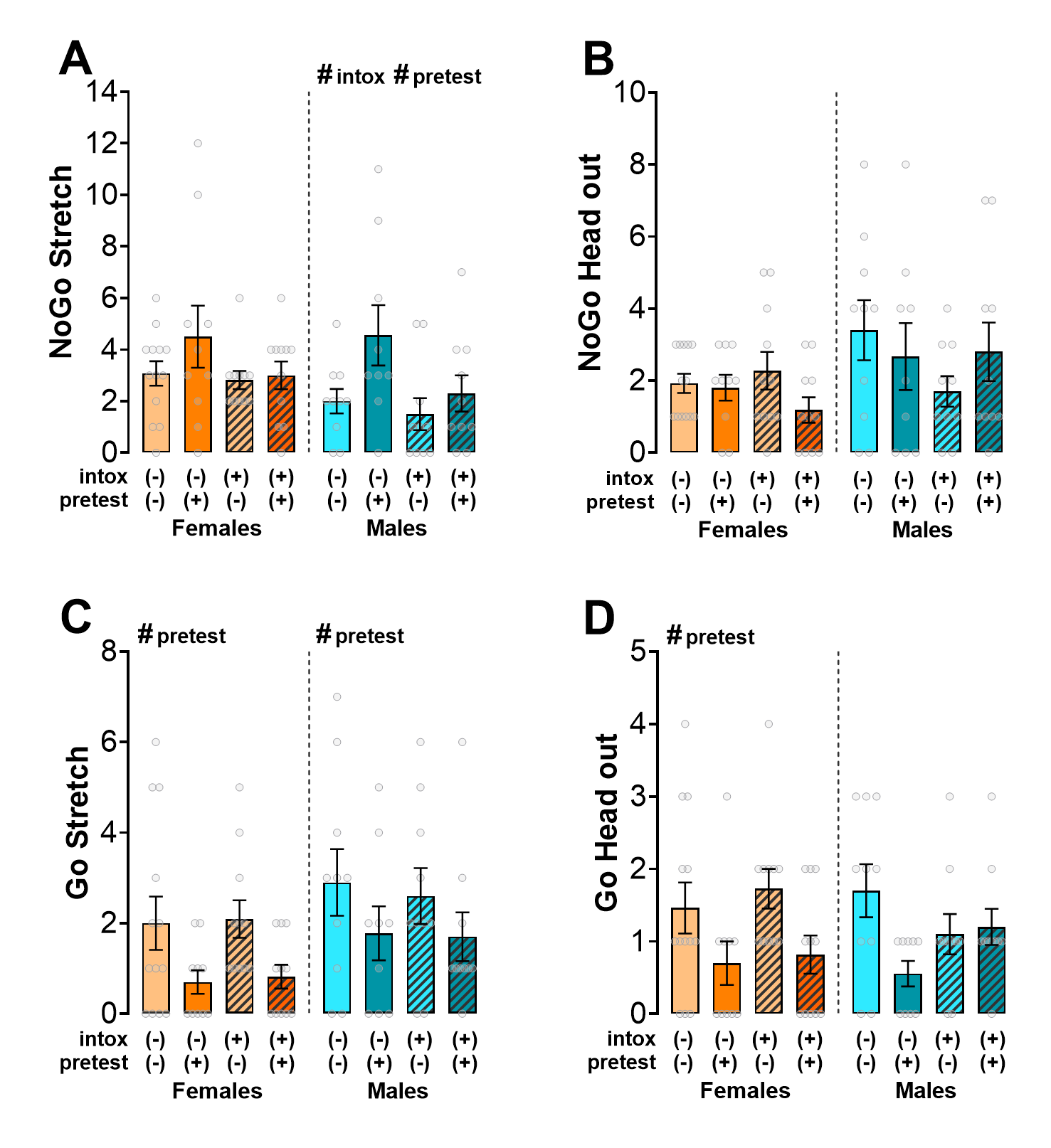

**Figure S2. NoGo and Go stretch-attend and head-out behaviors in the light–dark box (LDB) of Experiment 2.** All graphs show means ± standard error means (SEM). Orange-toned graphs represent females, and blue-toned graphs represent males. Hatched bars indicate intoxicated groups, which received ethanol during adolescence. Light-colored bars represent animals that received saline prior to testing in adulthood; dark-colored bars represent those that received ethanol prior to testing. Intox: intoxicated. **(A)** NoGo stretches. **(B)** NoGo head outs. **(C)** Go stretches. **(D)** Go head outs. #: p < 0.05 for main factor significance.

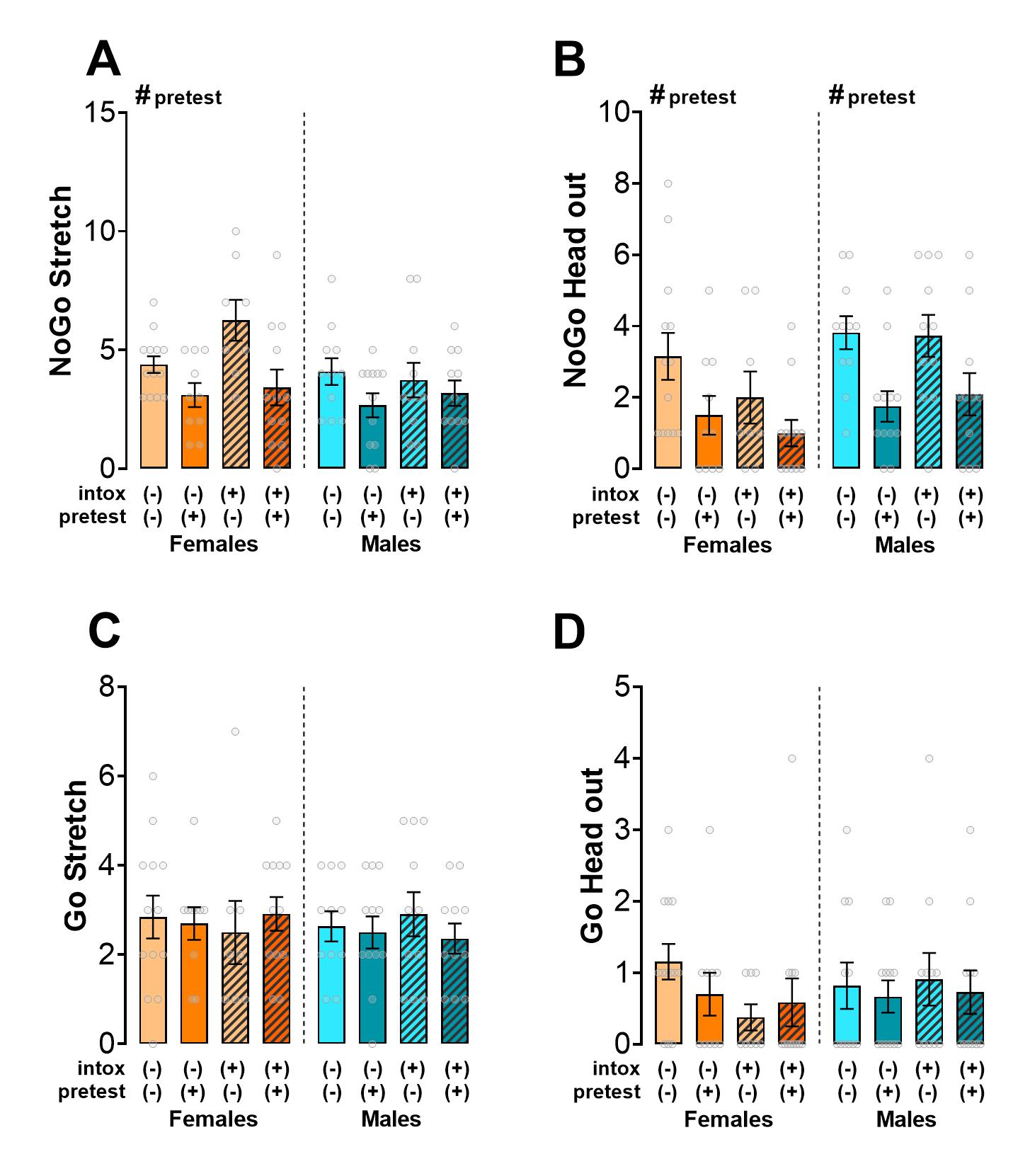

**Figure S3. NoGo and Go stretch-attend and head-out behaviors in the elevated plus maze (EPM) of Experiment 2.** All graphs show means ± standard error means (SEM). Orange-toned graphs represent females, and blue-toned graphs represent males. Hatched bars indicate intoxicated groups, which received ethanol during adolescence. Light-colored bars represent animals that received saline prior to testing in adulthood; dark-colored bars represent those that received ethanol prior to testing. **(A)** NoGo stretches. **(B)** NoGo head outs. **(C)** Go stretches. **(D)** Go head outs. #: p < 0.05 for main factor significance.

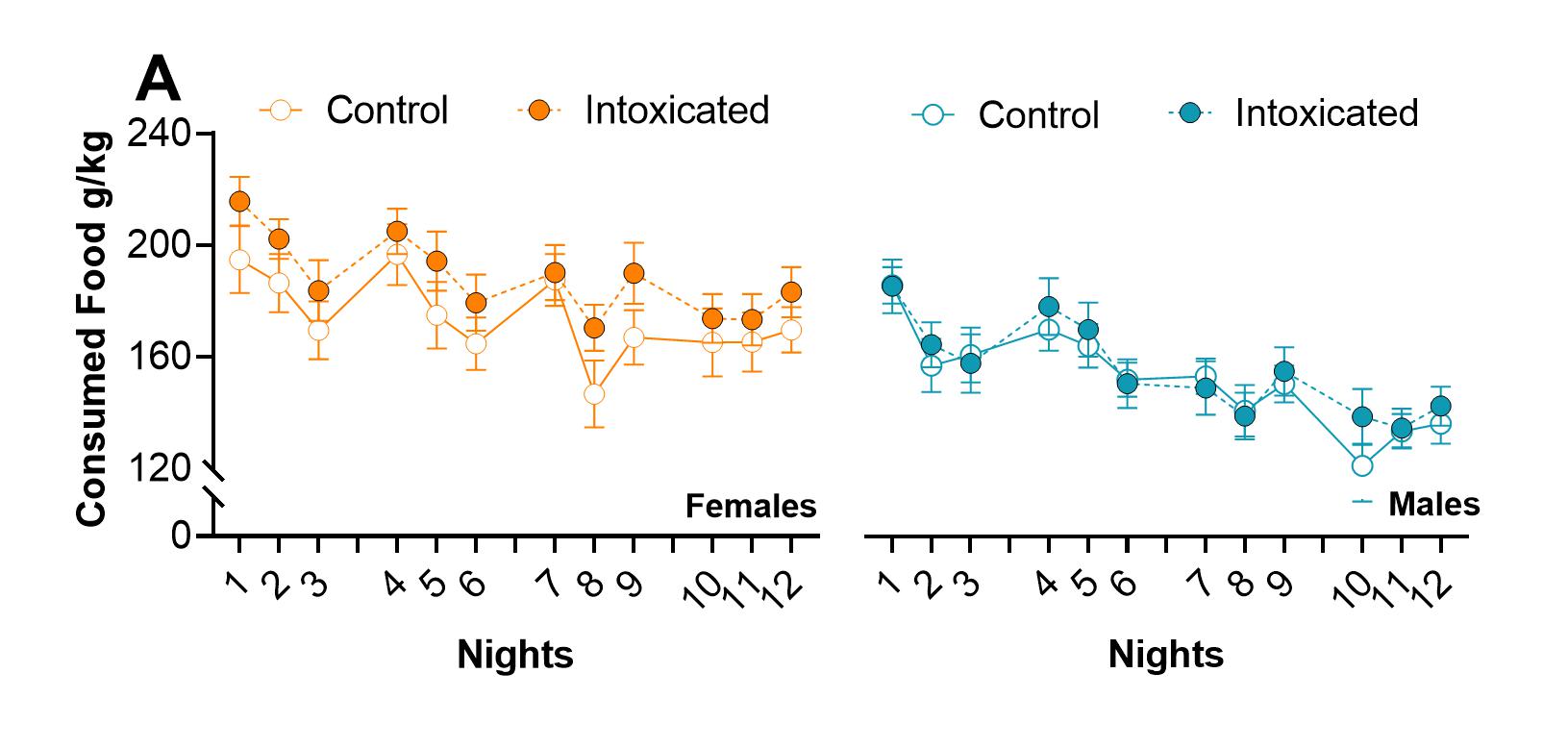

**Figure S4. Food intake during the IOD protocol of Experiment 3.** All graphs show means ± standard error means (SEM). Orange-toned graphs represent females, and blue-toned graphs represent males.
